# Maternal transfer of neutralizing antibodies to OspA after oral vaccination of the rodent reservoir

**DOI:** 10.1101/2021.01.27.428441

**Authors:** Kathryn O’Connell, Nisha Nair, Kamalika Samanta, Jose F. Azevedo, Grant D. Brown, Christine A. Petersen, Maria Gomes-Solecki

## Abstract

Lyme Disease presents unique challenges for public health efforts. We hypothesized that transfer of protective antibodies between mothers and offspring should occur after oral vaccination of C3H-HeN mice with *E. coli* overexpressing OspA. We present new evidence for maternal transfer of vaccine induced neutralizing anti-OspA IgG antibodies to mouse pups through ingestion of colostrum. Protective levels of OspA antibodies in pups were present from 2-5 weeks after birth and they persisted in some mice until 9 weeks of age. This was corroborated by detection of neutralizing antibodies in the serum of all pups at 2-3 weeks after birth and in some mice at 9 weeks of age. A clear association was found between robust antibody responses in mothers and the length of antibody persistence in the respective pups using a novel longitudinal Bayesian model. These factors are likely to impact the enzootic cycle of *B. burgdorferi* when reservoir targeted OspA-based vaccination interventions are implemented.

## Introduction

The global human and economic impact of Lyme borreliosis is vast, being especially prevalent throughout the northern hemisphere^1^, and to a lower degree in parts of Central and South America, Africa, and Asia^2^. Lyme borreliosis is caused by some species of the spirochete *Borrelia burgdorferi* sensu lato^3,4^, transmitted to mammalian hosts by *Ixodes* tick vectors^5^. The pathogen is maintained in the environment by tick-mediated infection of reservoir host species^6^, mostly rodents, rather than through transovarial transmission between tick cohorts^7^ or transmission between mammalian hosts^8^. In North America, the primary reservoir host species of *B. burgdorferi* is the white-footed mouse (*Peromyscus leucopus*)^6,9^.

Antibodies to outer surface protein A (OspA) protected mice from infection with *B. burgdorferi* by agglutination and neutralization of this spirochete^10^ within the tick midgut^11^ which leads to blockage of transmission. We developed a reservoir targeted vaccine based in *E. coli* overexpressing OspA and showed that oral administration of such vaccines to mice induced OspA-specific IgG antibodies in blood that prevented transmission of *B. burgdorferi* from the tick vector to the murine host. This was further complemented by drastic reductions of *B. burgdorferi* from *Ixodes* nymphs used for challenge^12^. White-footed mice (*P. leucopus*) vaccinated orally with baits containing *E. coli* overexpressing OspA maintained sufficient levels of neutralizing OspA-specific IgG in blood for up to a year^13^. Presence of protective levels of anti-OspA antibody in white-footed mice correlated with reductions in nymphal infection prevalence over time in a 5-year field trial^14^.

Transfer of antibodies from mother to offspring is a classical mechanism^15^ by which newborns are protected from many infectious diseases. To date there is no evidence of this type of protection against *Borrelia burgdorferi* infection. We hypothesized that maternal transfer of neutralizing anti-OspA antibodies occurred after vaccination of female mice. We immunized mice orally with the same vaccine master stock previously tested^12,13^, and evaluated transfer of OspA-specific antibodies to offspring via transplacental and transmammary transmission. Furthermore, we developed a novel Bayesian longitudinal model to jointly evaluate mother and pup immunity, allowing a robust exploration of the potential mechanisms for maternal transfer of antibody based on the time at which we detect antibodies in pups paired with the mother’s immune state.

## Methods

### Ethics statement

This study was carried out in accordance with the Guide for the Care and Use of Laboratory Animals of the US National Institutes of Health. The protocol was approved by the University of Tennessee Health Science Center (UTHSC) Institutional Animal Care and Use Committee, Animal Care Protocol Application Permit Number 19-0103.

### Animals

C3H-HeN mice were purchased from Charles River Laboratories and acclimatized for one week at the specific pathogen free (SPF) environment in the LACU-UTHSC.

### Bacterial strains

Glycerol stocks of *E. coli* BL21(DE3), *E. coli* BL21(DE3) transformed with pET9c-*ospA* and *B. burgdorferi* sensu stricto are kept at −80°C in the laboratory. Cultures of several strains of *B. burgdorferi* sensu stricto were recovered from heart and bladder tissues from white-footed mice naturally infected with *B. burgdorferi* (Bb) in 2005, 2006, 2007 and 2008. To generate a multi-strain culture for neutralization assays, 200ul of each stock was combined and grown to 1.00E07 Bb/mL in 7mL of Barbour-Stoenner-Kelly (BSK) media supplemented with 100x antibiotic mix for *Borrelia* and grown at 34°C for 2-4 weeks. *Borrelia* was enumerated by dark-field microscopy (Zeiss USA, Hawthorne, NY) and culture concentration was confirmed by qPCR using primers for *flaB* (StepOne Plus, Life Technologies, Grand Island, NY).

### Production of vaccine

Vaccine was produced according to modified published protocols^16^. Briefly, recombinant *E. coli* were grown in Tryptone Broth Yeast (TBY) medium supplemented with 50 μg/ml Kanamycin (Kn) at 37°C, shaking at 225 rpm, until it reached an OD_600_ of 0.8-0.9. The expression of recombinant OspA was induced by adding IPTG (isopropyl-β-d-thiogalactopyranoside, 0.5 - 1 mM final concentration) to the cells followed by incubation at 37°C, shaking 225 rpm, for 3 h. Non-transformed *E. coli* cells used as placebo control were grown in TBY without antibiotic. The cells were harvested by centrifugation at 4000 x g for 10 min at 4°C. Antigen expression was checked by Coomassie blue staining of constructs on polyacrylamide gels followed by immunoblot using anti-OspA monoclonal antibody 184.1. Aliquots of 1.00E10 cells/ml were resuspended in 10% glycerol-phosphate-buffered-salt solution and stored at −80°C. Aliquots were thawed at 4°C and ~1.00E09 *E. coli* cells (~300μl) were placed in a ball tipped disposable feeding needle (Fisher Scientific, Pittsburgh, PA) for oral gavage inoculation.

### Immunization schedule, breeding and collection of blood

Female mice were separated into 2 groups: one group of 10 females was assigned to the vaccine group and one group of 6 females was assigned to placebo. Females that received oral vaccine (EcA) were labeled as mothers M1 to M10 and females that received oral placebo (Ec, Ctrl) were labelled as mothers M11 to M16. A timeline diagram depicting vaccination, breeding and collection of samples is shown in **Fig. 1.**

Immunization schedule: 6-week old female mice received 1 vaccine dose (~10^9^ *E. coli* cells) daily by oral gavage for 2 full work-weeks (10 doses); mice rested for 1 week and on the following week they received the 1^st^ boost (1 dose/day for 5 days) and after resting another week they received a 2^nd^ boost (1 dose/day for 5 days) for a total of 20 vaccine doses per mouse.

**Fig. 1.**
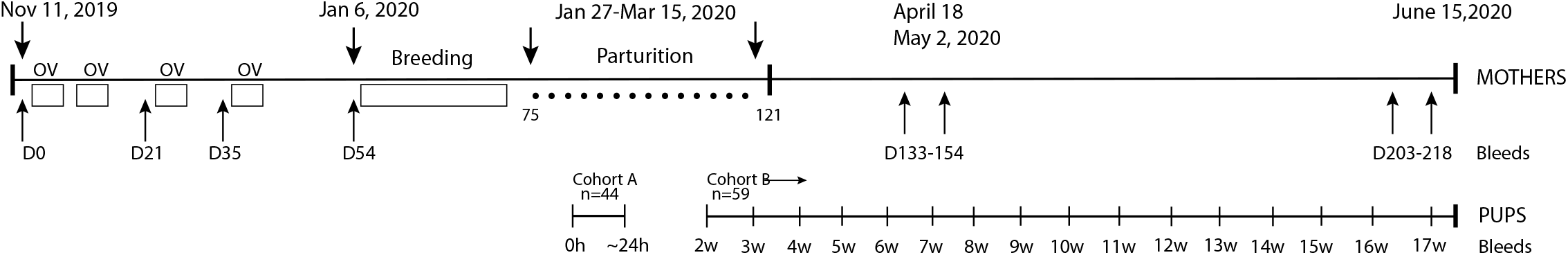
Diagram of the immunization schedule, breeding, parturition and bleeding of mothers and offspring. OV, oral vaccination; D, day after first vaccine dose; w, weeks.

Breeding and birth: 2 weeks after receiving the last vaccine dose (8-week vaccination schedule), pairs of females were co-housed with an age matched male for about 3-4 weeks. After breeding each female was single housed until parturition. Parturition occurred over a span of ~6 weeks due to differences in successful mating between pairs. Females started giving birth to pups ~11-17 weeks after the first vaccine dose. Pups were followed for 17 weeks.

Collection of blood for serology and neutralization assays: blood was collected from the mothers before immunization on D0, before the first boost on D21, before the second boost on D35, post-vaccination before breeding ~D54, post-vaccination post-birth on D133/154 (when the pups ~7 weeks old) and post-vaccination at termination ~ D203/218 (when the pups were ~17 weeks old). The pups were split into two cohorts. The first cohort of 44 pups (Cohort A) was sacrificed for collection of blood on the day of birth (most pups were born overnight). Cohort A included 12 control pups born to Mother 11 and from litters from Mothers 14 and 16, and 32 pups born to the first litter of vaccinated Mothers 6-10. The second cohort (n=59, Cohort B) was comprised of 33 pups born to the first litter of control Mothers 12-16, and 27 pups from the first litter of vaccinated Mothers 1-5; blood was collected via the submandibular vein on a weekly basis starting 2 weeks after birth until the pups reached 17 weeks of age.

### ELISA assay

ELISA was performed according to adapted published protocols^13,17^. Briefly, 1 μg/mL of purified OspA^18^ is used to coat flat-bottom MaxiSorp ELISA plates (eBioscience) overnight; next morning the plate is washed, blocked, incubated with the 1^st^ antibody (mouse serum diluted at 1:100) followed by the second antibody HRP conjugated goat anti-mouse IgG (Jackson ImmunoResearch) diluted at 1:10000, TMB substrate and stop solution according to steps 16-28 of Basic Protocol 3 in^17^. The plates were read in a SpectraMax ELISA reader (Molecular Devices) at 450nm.

### *B. burgdorferi* immunoblot

Multi-strain *B. burgdorferi* was cultured in BSK media at 34° until the cell count reached 10^8^/ml; 1 ml of the culture was centrifuged, washed in PBS, further diluted to 1:20 in PBS and 25μL 4X gel loading dye containing ß-Mercaptoethanol was added to 75 μL of the solution. The extract was boiled for 5 minutes at 95°C before SDS-PAGE was performed. The proteins were transferred onto PVDF by placing the gel on a Bio-Rad semi-dry transfer system. Western blot analysis was performed using a monoclonal antibody to OspA (mAb 184.1, 1:1000) and an alkaline phosphatase labelled secondary anti-antibody (1:5000).

### Neutralization assay

Presence of neutralizing anti-OspA antibody was determined in serum from mothers and pups as previously described^13^. We used serum from mothers collected after vaccination and before breeding (~D54). The serum from pups was collected on the day of birth (0-24h), at 2-3 weeks after birth and at 9 weeks after birth. Briefly, a 10^7^/mL multi-strain culture of *B. burgdorferi* was incubated with heat inactivated mouse serum in the presence of guinea pig complement and incubated for 6 days at 34°C; cultures were checked on days 3 and 6 and the number of viable, motile *B. burgdorferi* were counted in a Petroff-Hausser chamber under a dark field microscope.

### Bayesian Model

Rather than considering group differences and treating pup and mother analyses separately, we introduced a Bayesian model to jointly consider anti-OspA antibody levels in mothers and her pups. Specifically, we introduced a pair of auto-correlated, log-scale time-series with forcing parameters related to oral vaccine dose-timing (mothers) and birth/weaning processes (pups). Parameters were estimated separately between vaccine and placebo groups. For experimental group *i*, mother *j*, and time index *t*, we modeled the outcome as an auto-regressive Gaussian time series (Eq. 1).

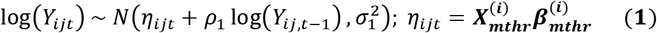

The design matrix ***X***^(***i***)^ contains an intercept, a linear-time dose term (up to D54), and a linear-time change point for decay (starting at D54). Pups are coupled to these series by matching them to their respective mothers. For vaccinated group *i*, pup *j*, and time *t*, we defined (Eq. 2):

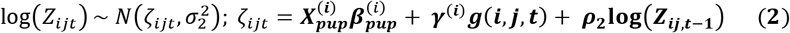

Let *g*(*i, j, t*) be zero, except on the first pup-measurement for pups not sacrificed immediately post-parturition, at which point it is the matched maternal D54 anti-OspA antibody level in blood. This set up permitted the model to directly investigate evidence for transplacental antibody transfer by allowing separate intercepts by treatment group, while simultaneously investigating the potential for post parturition transfer by coupling the pup anti-OspA level in blood to matched mothers. Vague priors were specified:

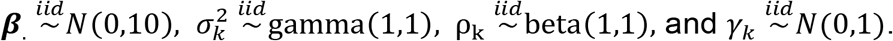

The model was implemented as a reproducible R-markdown document and is referenced along with the requisite data sets in **Table S1**. The analysis presents a novel approach to joint modeling of mechanistically-related longitudinal data sets and allows us to make full use of the data from complex experiments and directly measure the effects of interest, rather than relying on informal comparisons or more basic significance tests. This approach can be extended and generalized, allowing direct application to follow-up studies. Furthermore, the model can be applied to other microbiological problems with dependent longitudinal relationships. The model was fit using the `rstan` R package interface to the `Stan` statistical software in R version 4.0.2^19, 20^.

## Results

A total of 105 C3H-HeN pups were born to the 16 mothers: 59 pups were born to mothers vaccinated with *E. coli* overexpressing OspA (EcA), and 46 pups were born to mothers that received *E. coli* placebo control (Ec, Ctrl) with an average litter size of 5.9 for EcA and 5.75 for Ctrl. Insufficient blood was collected from 2 pups born to control mothers, and as such, those pups were not included in further analysis.

### Kinetics of αOspA IgG in serum from orally vaccinated mothers over 7 months

We measured anti-OspA IgG in serum from orally vaccinated C3H-HeN mothers and placebo controls by ELISA (**Fig. 2**). As expected, the prevalence of mice with protective levels of OspA antibody, defined by OD_450_ ≥ 0.8, increased in EcA vaccinated mothers from 20% at 21 days after receiving the first vaccine dose, to 60% after the first boost (D35), to 90% after the second boost (D54). Five months after mothers received their first oral vaccine dose (~D133/154), prevalence of protective levels of antibody was 77.8%. Two months later, 7 months after the first dose ~ D203/218, prevalence of protective anti-OspA antibody was 30%. Controls that received *E. coli* placebo (Ec) did not develop OspA-specific IgG. Differences between EcA and Ec groups were statistically significant throughout the vaccination schedule (Fig. 2A). Of the 10 mothers that received the oral vaccine by gavage, Mothers 3, 4, 5, 6 and 9 developed the most robust IgG response to OspA on D54 (Fig 2B).

**Fig. 2.**
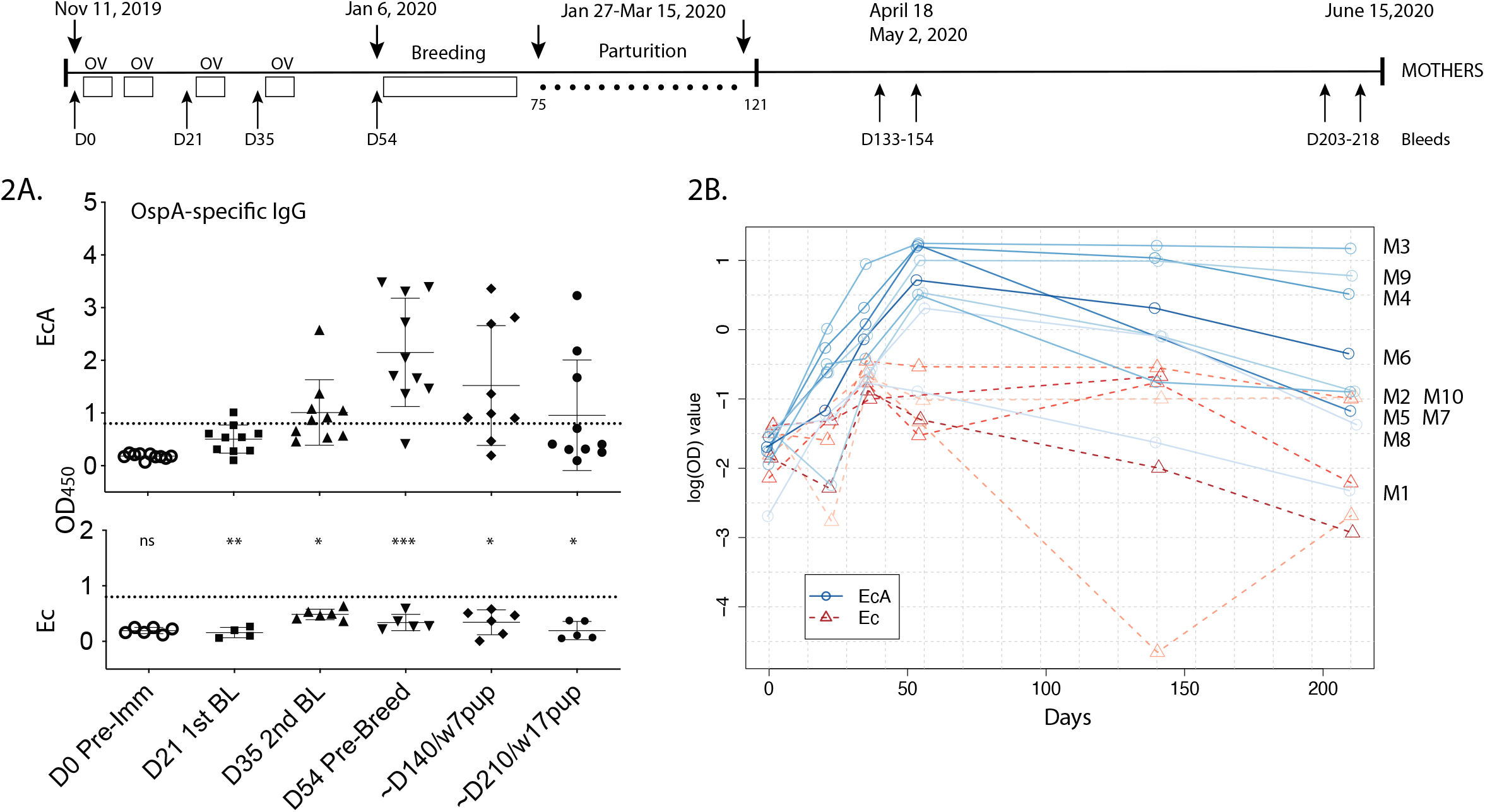
Kinetics of OspA-specific antibodies in orally vaccinated and placebo mothers over ~7 months. We measured anti-OspA IgG by ELISA in serum from 10 mothers orally vaccinated *E. coli* expressing OspA (EcA, M1-M10) and 6 non-vaccinated placebo controls (Ec, M11-M16). Mice were bled before (D0) and after immunization (D21, D35), before breeding (D54) after the pups were born ~D140 (7 wk old pups) and ~D210 (17 wk old pups). A, Comparison of anti-OspA IgG (OD450>0.8) between EcA and placebo Ec groups. Differences between EcA and Ec groups are statistically significant by unpaired t test with Welch’s correction, D21 p=0.0038, D35 p= 0.0260, D54 p=0.0003, D140 p=0.0147, D210 p=0.0488. B, Longitudinal log scale trajectory of antibody response over time, each line represents one mother, M5 was not bled on D140.

### OspA-specific IgG was detected in serum from pups born to vaccinated mothers

We measured anti-OspA IgG in serum from offspring born to the vaccinated mothers from two cohorts of pups: one cohort was terminated on the day of birth to access transplacental transfer of maternal antibodies and a second cohort of pups was bled over a period of 4 months, weekly from week 2-17 after birth, to access transfer of transmammary antibodies through colostrum and milk (**Fig. 3**). A longitudinal analysis of anti-OspA IgG between EcA and Ctrl mice taken at all time points showed a clear distinction between the two groups, except for the pups terminated on the day of birth (Fig 3A). In the cohort of pups terminated 0-24h after birth (Cohort A), overall differences in OspA-specific antibody were not significant between pups born to EcA and Ctrl mothers. However, a breakdown of data by birth mother showed there was one litter of pups born to M9 that clearly had anti-OspA antibodies in serum (Fig. 3B). In the second cohort of mice (Cohort B) we observed an increased anti-OspA IgG mean at 2 weeks post birth (OD_450_=1.649, 95% CI 0.677 to 1.832) that steadily decreased until 9 weeks post birth (OD_450_=0.382, 95% CI 0.019 to 0.366). Differences are significant from 2-9 weeks after birth: 2w p=0.0002, 3w p=0.0001, 4w p=0.0010, 5w p=0.0006, 6w p=0.0024, 7w p=0.0249, 8w p=0.0144 and 9w p=0.0309. When we analyzed the anti-OspA IgG by birth mother we observed significant differences in protective levels of anti-OspA IgG (OD_450_>0.8) between litters born to 4 of 5 mothers (Fig. 3C). Pups born to mother M1, which never developed anti-OspA antibodies, did not have antibodies in serum. Interestingly, pups from larger litters had less total antibody to OspA in serum and pups with the highest levels of antibodies to OspA were born to the mothers of this cohort that produced the most robust response to OspA, M3, M4, M5 (Fig. 3C), which suggested a correlation between these two parameters.

**Fig. 3.**
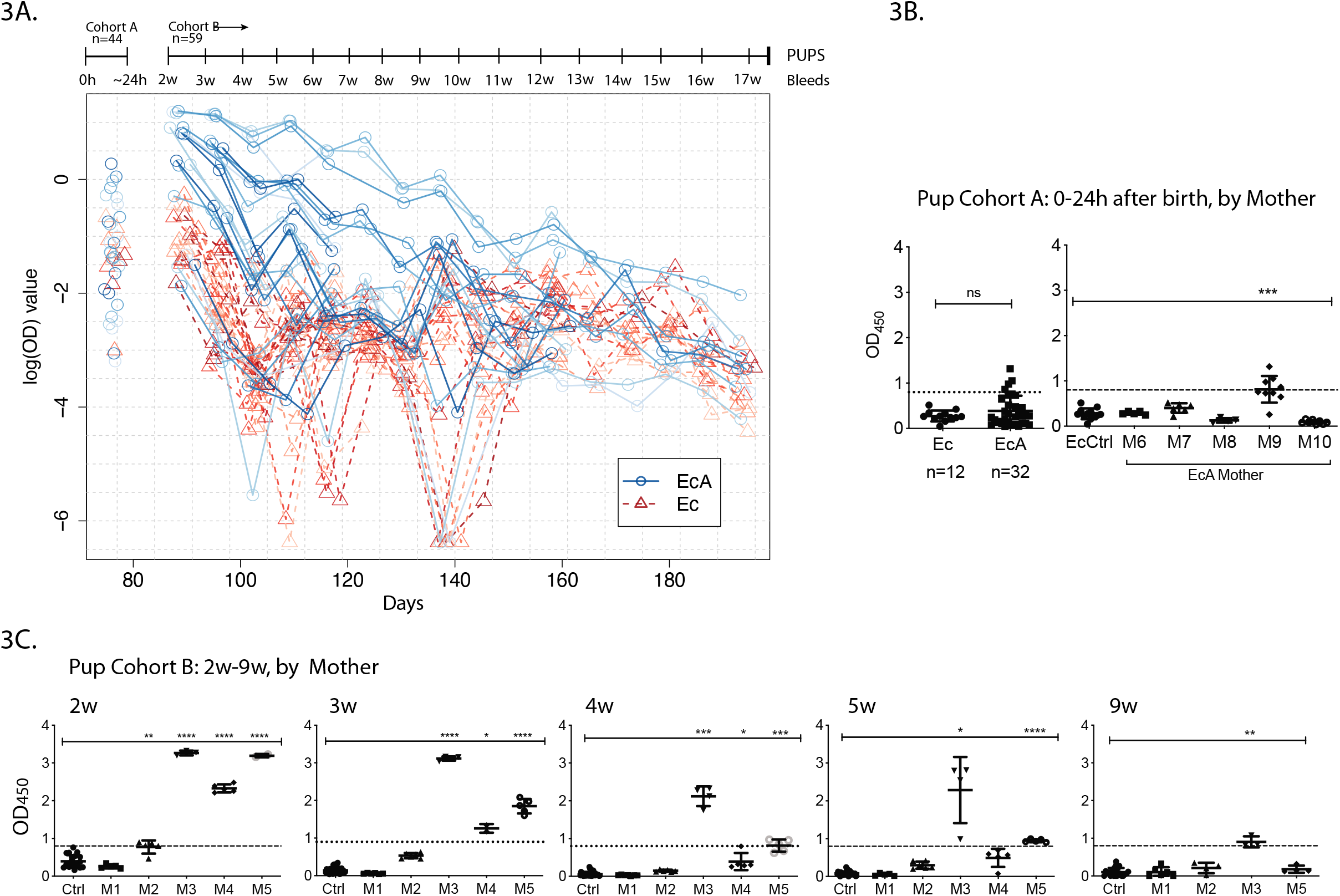
Kinetics of OspA-specific antibodies in offspring born to orally vaccinated mothers at birth and over 4 months post birth. A, Longitudinal log scale trajectory of anti-OspA IgG measured in serum from both cohorts of pups: Cohort A pups analysed on the day of birth, and Cohort B pups analysed 2 to 9 weeks after birth. B and C, Anti-OspA IgG in both cohorts of pups grouped by mother. Statistics by the Unpaired t test with Welch’s correction between EcA and Ec Ctrl, * p<0.05, ** p<0.005, *** p<0.0005 and **** p<0.0001. EcA, oral vaccination with E. coli expressing OspA; Ec Ctrl, placebo control; M1-M10 EcA vaccinated mothers; w, weeks.

### Maternal and pup antibody levels unambiguously connected

To further investigate the correlation between amount of anti-OspA maternal antibody and the amount of antibody transferred to their respective offspring we modelled these two components of maternal/fetal antibody transfer in a Bayesian Hierarchical Model framework. We found clear evidence of a treatment effect in mothers and a strong correlation between the amount of protective antibody in vaccinated mothers and their respective offspring. Due to non-specific antibody activity, both EcA and control mothers exhibit evidence of a positive dose effect, with probabilities of an effect greater than zero: ≈ 1 in the EcA group and 0.97 in the control group. Antibody trends over time were very different between EcA and Ec, since all parameters like intercepts, dose, decay, autocorrelation, were estimated separately (**Table 1**). We estimate a 99% probability that the effect of maternal vaccination on presence of anti-OspA antibodies in the pups was higher in EcA group than Ec (**Fig. 4**). In addition, the EcA group exhibited higher autocorrelation (probability 0.88). For pups, the treatment effect is primarily seen through shifts in the intercept, capturing birth or shortly post-birth antibody activity and the D54 mother OspA-specific antibody capturing exposure post-parturition. Pups exhibited an elevated baseline antibody level relative to controls (posterior probability ≈ 1), and also demonstrated a positive and elevated D54 weaning term (posterior probabilities both ≈ 1). Crucially, Ec pups exhibited significant evidence (probability 0.99) of a small, negative weaning effect. Thus, despite the presence of nonzero antibody activity in both groups, we observe unambiguous differences between vaccine-induced OspA-specific antibodies and unmodified cross-reactive antibodies in the control group. In addition, we find evidence (probability ≈ 1) that autocorrelation was higher in EcA group pups than in control group pups, and that the decay of antibody levels was slower (probability 0.85).

**Figure 4.**
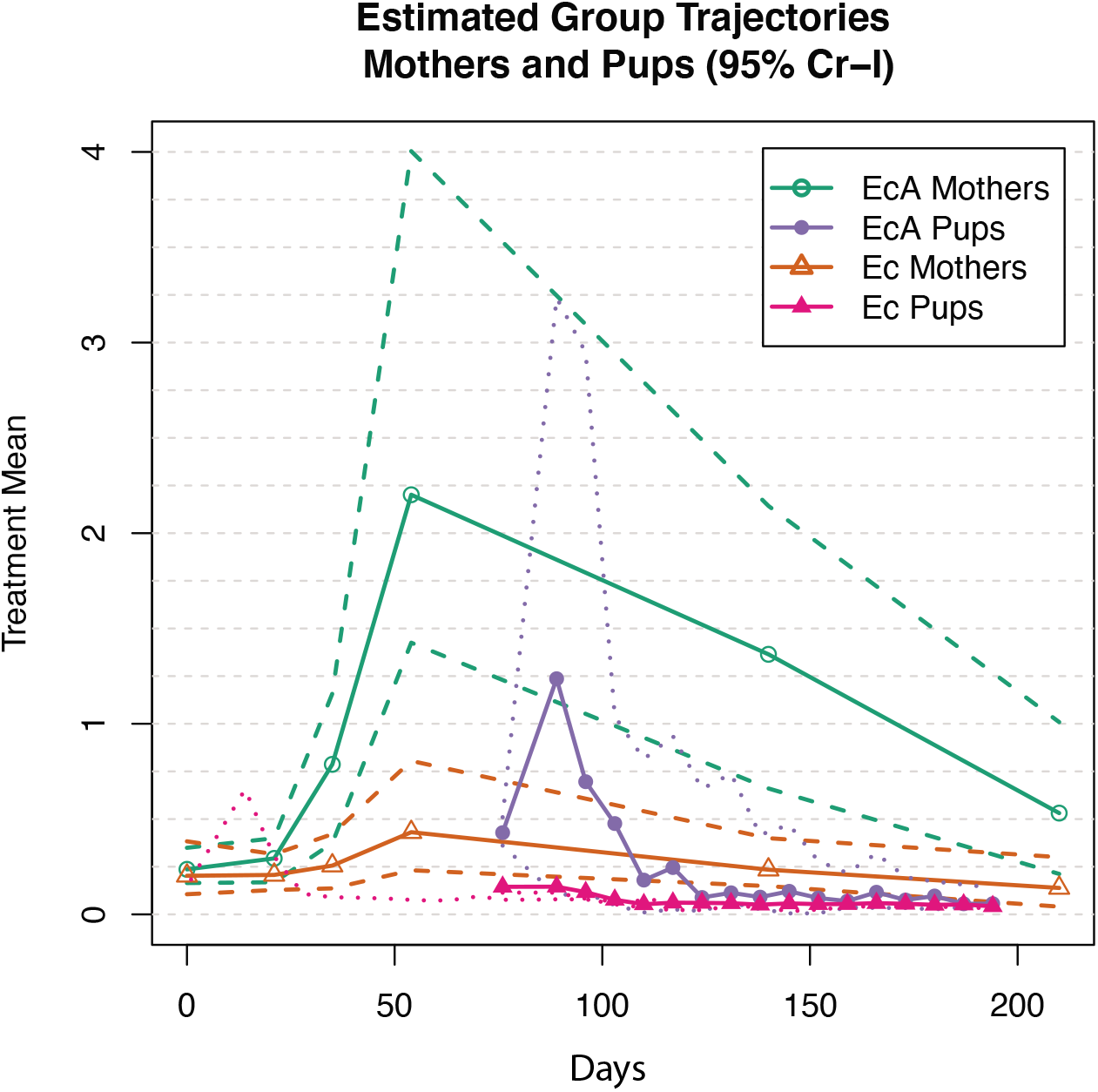
Bayesian hierachical model showing median anti-OspA antibody levels in EcA vaccinated mothers, control mothers, and pups from each group. Each group median is given along with 95% posterior credible intervals (Cr-I). Differences in antibody curves between vaccinated and unvaccinated mothers and their offspring are immediately apparent. Legend: EcA, oral vaccination with E. coli expressing OspA; Ec, oral vaccination with Ec placebo; Cr-I, Credible Intervals.

**Table 1:** Parameter Estimates and Comparisons

| Table 1: Parameter Estimates and Comparisons |  |  |  |  |
| --- | --- | --- | --- | --- |
| Parameter Estimates |  |  |  |  |
| Parameter | Cohort | Mean (95% CrI) | P(x>0) | P(x<0) |
| **EcA** Intercept | Mothers | -1.44 (-1.81, -1.05) | 0 | 1 |
| **EcA** Linear Dose Effect | Mothers | 2.38 (1.82, 2.93) | 1 | 0 |
| **EcA** Decay Term | Mothers | -3.9 (-5.21, -2.58) | 0 | 1 |
| Control Intercept | Mothers | -1.6 (-2.25, -0.96) | 0 | 1 |
| Control Linear Dose Effect | Mothers | 0.94 (-0.08, 1.96) | 0.97 | 0.03 |
| Control Decay Term | Mothers | -2.48 (-4.88, -0.07) | 0.02 | 0.98 |
| **EcA** Autocorrelation | Mothers | -0.85 (-1.02, -0.67) | 0 | 1 |
| Control Autocorrelation | Mothers | 0.03 (-0.4, 0.46) | 0.56 | 0.44 |
| **EcA** Intercept | Pups | -1.94 (-2.17, -1.7) | 0 | 1 |
| **EcA** Decay Term | Pups | -0.34 (-0.78, 0.09) | 0.06 | 0.94 |
| Control Intercept | Pups | 0.4 (0.2, 0.6) | 1 | 0 |
| Control Decay Term | Pups | 0.2 (0.01, 0.5) | 1 | 0 |
| **EcA** Autocorrelation | Pups | 0.69 (0.61, 0.77) | 1 | 0 |
| Control Autocorrelation | Pups | 0.27 (0.19, 0.36) | 1 | 0 |
| **EcA** Pup Maternal Exposure Term | Mothers/Pups | 1.39 (1.08, 1.71) | 1 | 0 |
| Control Pup Maternal Exposure Term | Mothers/Pups | -0.01 (-0.02, 0) | 0.01 | 0.99 |
| EcA vs. Control (Ec) Comparisons |  |  |  |  |
| Comparison | Cohort | Mean (95% CrI) | P(x>0) | P(x<0) |
| Intercept | Mothers | 0.16 (-0.59, 0.91) | 0.67 | 0.33 |
| Linear Dose Effect | Mothers | 1.44 (0.26, 2.6) | 0.99 | 0.01 |
| Decay | Mothers | -1.42 (-4.16, 1.35) | 0.15 | 0.85 |
| Autocorrelation | Mothers | 1.09 (0.79, 1.39) | 1 | 0 |
| Intercept | Pups | 0.37 (-0.24, 0.98) | 0.89 | 0.11 |
| Decay | Pups | 0.2 (-0.14, 0.48) | 0.88 | 0.12 |
| Autocorrelation | Pups | 0.42 (0.3, 0.54) | 1 | 0 |
| Maternal Exposure Term | Mothers/Pups | 1.41 (1.09, 1.72) | 1 | 0 |

### Serum from vaccinated mothers and their pups neutralize *B. burgdorferi* in culture

The function of neutralizing antibody (nAb) was analyzed in serum from EcA and Ctrl mothers before the females were bred on ~D54, and on the blood of their respective offspring (**Fig. 5**). nAb was measured by dark field microscopy quantification of motile *B. burgdorferi* in multi-strain cultures treated with serum from vaccinated and placebo mothers over two time points. Due to insufficient sample to test all mothers independently, we pooled serum from the immune mothers used for the analysis of long-term kinetics of antibody to OspA (M2, M4, M5) and from the mothers used for analysis of antibody transfer at birth (M6, M7, M8, M10). All the mothers included in this analysis (M2 to M10) had levels of OspA-specific antibodies OD_450_>0.8 before breeding on D54 (Fig. 2A). We did not include M1 because this mother did not develop antibodies to OspA. We were able to keep one mother from each group used to generate the two cohorts of mice (A and B) that maintained highest levels of antibody to OspA on ~D210 (M3 and M9), to test independently. We ran an immunoblot of a multi-strain culture used for the neutralization assay against an anti-OspA monoclonal antibody and show that OspA is expressed by *B. burgdorferi* (Bb) in culture (**Fig S1**). We found that serum from mothers M6 to M10 that produced pups used to evaluate transplacental antibody transfer on the day of birth (Cohort A), had sufficient anti-OspA antibody to reduce the number of motile Bb in culture by about 1Log_10_ (Fig 5A); serum from the mothers that produced Cohort B pups (pooled M2, M4, M5, and individual serum from M3) used to evaluate transfer of maternal antibody through colostrum and milk, had sufficient anti-OspA antibody to reduce the number of motile Bb in culture by about 7Log_10_ (Fig 5A). Serum from M9 pups collected on the day of birth did not have enough anti-OspA antibody to neutralize motile Bb in culture (Fig 5B). Serum from 2-3 week old pups born to M2-5 all had enough anti-OspA antibody to neutralize motile Bb in culture by 1-2 Log_10_ (M2, M3 and M5) and by 7 Log_10_ (M4) (Fig 5C). Serum from pups born to M3 still had enough anti-OspA antibody to neutralize motile Bb in culture by 1 Log_10_ 9 weeks after birth (Fig 5F).

**Figure 5.**
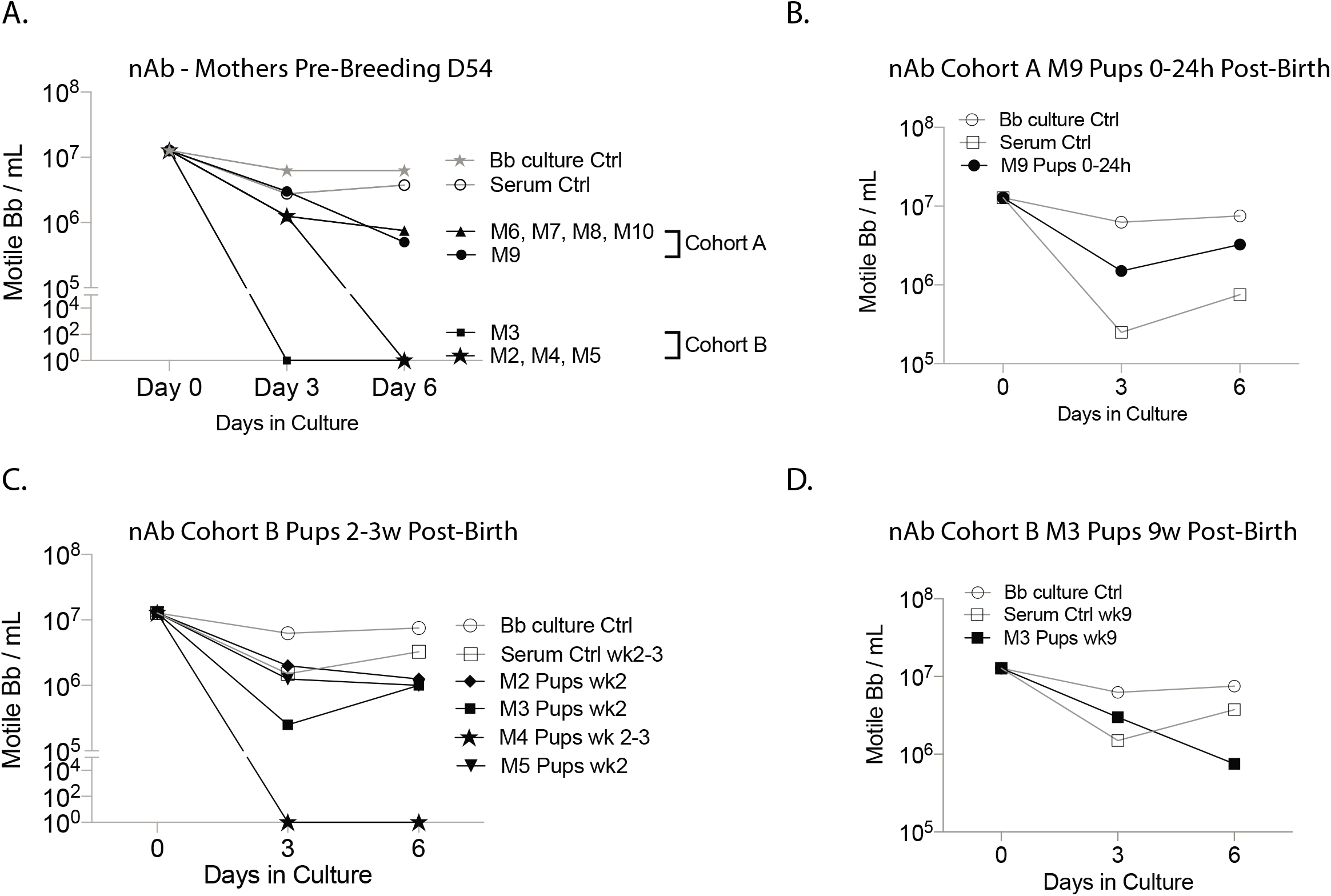
Anti-OspA neutralizing antibody (nAb) function against *B. burgdorferi*. Number of motile *B. burgdorferi* (Bb) after treatment of a multistrain culture with A) pooled serum from mothers M6 M7 M8 M10, pooled serum from mothers M2 M4 M5 and individual serum samples from mothers M3 and M9; B, pooled serum from all pups born to M9 on the day of birth (Cohort A); C, pooled 2-3w serum from Cohort B pups born to M2 M3, M4 and M5 (serum was pooled according to birth mother); and D, pooled serum from all pups born to M3 collected 9 weeks after birth (Cohort B). Live Bb were counted under a dark field microscope.

## Discussion

In this study we demonstrate that specific antibodies from immunized mouse mothers were transferred to offspring born to mice orally vaccinated with *E. coli* overexpressing OspA. Immunoglobulin G specific to OspA was passed via transmammary transmission to pups through ingestion of colostrum and milk during the lactation period, shortly after birth lasting 3 weeks. Although likely, we were unable to conclude with all confidence that antibodies are not transferred via the placenta. We found a strong statistical correlation of antibody transfer between mothers that produced the most robust IgG response to OspA and their respective pups. Presence of protective levels of antibodies lasted until 4-5 weeks of age in the majority of pups but persisted in some until 9 weeks. We also showed that serum obtained from vaccinated mothers and from respective offspring neutralized motility of cultured *B. burgdorferi* containing several spirochete strains demonstrating antibody function as a correlate of protection.

*B. burgdorferi* is maintained in the environment by tick-mediated infection of reservoir host species^6^, mostly rodents, rather than through transovarial transmission between tick cohorts^7^ and between mammalian hosts^8^. As a logical follow up to our previous findings we hypothesized that maternal transfer of vaccine-induced OspA-specific IgG to offspring would explain how the enzootic cycle of *B. burgdorferi* can be disrupted by oral bait immunization of field mice with OspA vaccines^21 14^. We vaccinated C3H-HeN mothers using a standard intermittent 8-week immunization schedule for oral gavage of an *E. coli* culture overexpressing OspA and followed the IgG response for 5 months in mothers and pups. We found that maternal oral vaccination led to an exponential increase of anti-OspA IgG three weeks post first vaccine dose. This started to decline after eight weeks. This is consistent with our previous findings with the same vaccine formulation and schedule^12^.

In the current study, we extended the schedule of blood collection until day 203-218 post first vaccine dose due to the unforeseen complication of animal work during the COVID-19 pandemic. We found that the mean of anti-OspA IgG remained above OD_450_ ~0.8 for about ~ 5.5 months after the 1^st^ boost (D35) in C3H-HeN mice vaccinated by oral gavage with a live culture of *E. coli* overexpressing OspA (Fig 2A). In our previous studies^13,14^ we determined that, unlike levels of LA-2 equivalent antibodies, an OspA ELISA 2<OD_450_>0.8 can be used to empirically establish the minimum levels of protective/neutralizing antibodies in serum from adult mice orally vaccinated with *E. coli* expressing OspA. We found that the number of mice with protective levels^13^ of anti-OspA IgG remained above 60% for about ~3.5 months (between D35 until D154), and it remained above 30% for two additional months. This is important as vaccination of foxes with an oral baited vaccine showed that 30% vaccine antibody prevalence in immunized foxes resulted in disrupted transmission of Rabies virus^22^. In addition, the strength of anti-OspA immunity differed considerably between the mothers (Fig 2B). This is also consistent with our previous findings using the same vaccine formulation and schedule^12^. Nevertheless, vaccination of *P. leucopus* using a baited formula rather than oral gavage and a different schedule of immunization^13^ it is reasonable to expect that oral bait administration of this vaccine should result in a longer and more robust immune responses to OspA.

Transfer of immune elements from mother to offspring via placenta and milk is a classical mechanism by which neonates are protected from infections acknowledged for over 120 years^15,23,24^. When we tested for presence of anti-OspA antibodies in pups born to vaccinated mothers we found a clear difference in antibody levels at birth versus 2+ weeks after birth (Fig 3). After birth, antibodies to OspA remained detectable in the pups for about 2 months. Furthermore, the robustness of the maternal immune response to OspA directly correlated with the amount of antibody transferred to offspring. We observed a positive association between high levels of antibodies in the birth mother and the length of time these antibodies persisted in the offspring serum; for example, the mother with the highest level of anti-OspA antibodies produced offspring that had persistent antibodies 9 weeks after birth, indicating a more durable antibody response. Another interesting finding was that pups from larger litters had less total antibody to OspA in serum which strongly suggests a dilution effect between mother and offspring. In other words, the mother produces a fixed amount of antibody to OspA at the peak of her immune response and this is split between the offspring depending on time of suckling. At birth, the pups born to 4/5 mothers did not have antibody to OspA. However, one mother (M9) produced pups that had antibodies to OspA on the day of birth. Since most of the pups were born overnight, M9 may have been the first mother to give birth and her pups may have begun nursing much sooner than the pups born to the other mothers. Although the data is highly suggestive of absence of transplacental transfer, we cannot reach an unequivocal conclusion, given that the mothers that produced the Cohort A pups also developed less overall antibody to OspA after vaccination. This difference could be explained by the longer time these mothers took to breed and produce offspring.

Dual modelling of mothers and their corresponding pups, we observed systematic differences in antibody profiles. In addition to the unambiguous detection of immunization effects in mothers and the presence of maternal transfer of antibodies to pups (Fig 4), we validated that maintenance of antibody activity was substantially different between vaccine and placebo groups. In addition, these comparisons remained clear in the presence of substantial effect heterogeneity from both mothers and pups, arising from factors such as the timing of birth of some litters, variability in maternal antibody levels, litter size variation, and numerous other factors. In addition to providing an efficient use of all the data available for these paired cohorts, this approach provides a template for how similar studies might make use of Bayesian hierarchical models. These techniques allow researchers to directly explore mechanisms under study and can be of substantial use in a wide variety of problems where complex dependence among data sets is expected.

The protective mechanism of action of OspA based vaccines relies on the OspA antibody ability to neutralize motile spirochetes in the nymphal tick midgut, thus preventing migration to the tick salivary gland and subsequent transmission to the next mammalian host^11^. When feeding on an OspA vaccinated mouse, an infected tick ingests a bloodmeal loaded with OspA antibodies that neutralize *B. burgdorferi* before it can be transmitted to the host. We and others have developed transmission blocking vaccines based in OspA^12,21,25^. Our reservoir targeted vaccine reduced prevalence of infected nymphal ticks by 76% after vaccination of the primary reservoir host species (*P. leucopus*) in a 5-year field trial^14^. In the study reported here we show that 90% of vaccinated mothers produced neutralizing anti-OspA antibodies for 4-5 months after receiving the last vaccine dose. Transfer of these antibodies from mother to offspring happened through consumption of colostrum and milk and neutralizing antibodies persisted in pups through two months of age (Fig. 5). We speculate that in a field setting where adult female mice are vaccinated in April and May, transfer of neutralizing anti-OspA antibodies from mother to offspring over a 4-5 month period through late Spring and Summer may lead to a population of reservoir offspring with significant levels of neutralizing anti-OspA antibodies in blood through July and August when the uninfected larva start to feed. This factor is likely to impact the enzootic cycle of *B. burgdorferi*. This novel observation may explain how vaccinating field mice through the summer with baits containing *E. coli* expressing OspA lead to maintenance of sufficient neutralizing anti-OspA antibodies in the offspring to prevent new reservoir host infection into the larval feeding season and may have reduced acquisition of *B. burgdorferi* by the next nymphal tick cohort the following spring^14^. Further experimentation will shed light on maternal transfer of anti-OspA antibody on larval and nymphal infection prevalence and will allow us to make unambiguous conclusions regarding transplacental transfer of anti-OspA antibodies.

## Acknowledgements

This work was supported by the Public Health Service awards AI139267 (to MGS and CAP) and AI155211 (to MGS) from the National Institute of Allergy and Infectious Diseases (NIAID) of the National Institutes of Health (NIH) of the United States of America. The content of this manuscript is solely the responsibility of the authors and does not necessarily represent the official views of NIAID or NIH. Kamalika Samanta is supported in part by the Doctoral Program in Pharmaceutical Sciences at UTHSC.

## Author contributions

Conceptualization and supervision: Maria Gomes-Solecki

Bayesian model development, implementation, and analysis: Grant Brown

Experimental investigation: Kathryn O’Connell, Nisha Nair, Kamalika Samanta, Jose F. Azevedo

Data analysis: all authors

Funding acquisition: Maria Gomes-Solecki and Christine Petersen

Writing – original draft: Maria Gomes-Solecki; and Bayesian model, Grant Brown

Writing – review and editing: all authors

## Competing interests

The following authors declare potential conflicts of interest: MGS (grants from federal agencies, employment, patents and consultant), CAP (grants from federal agencies, employment, and consultant). All other authors declare no conflicts.

