## Supplemental material for "Maternal transfer of neutralizing antibodies to OspA after oral vaccination of the rodent reservoir"

Table S1.

Full code and data to reproduce the Bayesian model is available at the following link:
https://github.com/grantbrown/Lyme-RTV-Maternal-Transfer

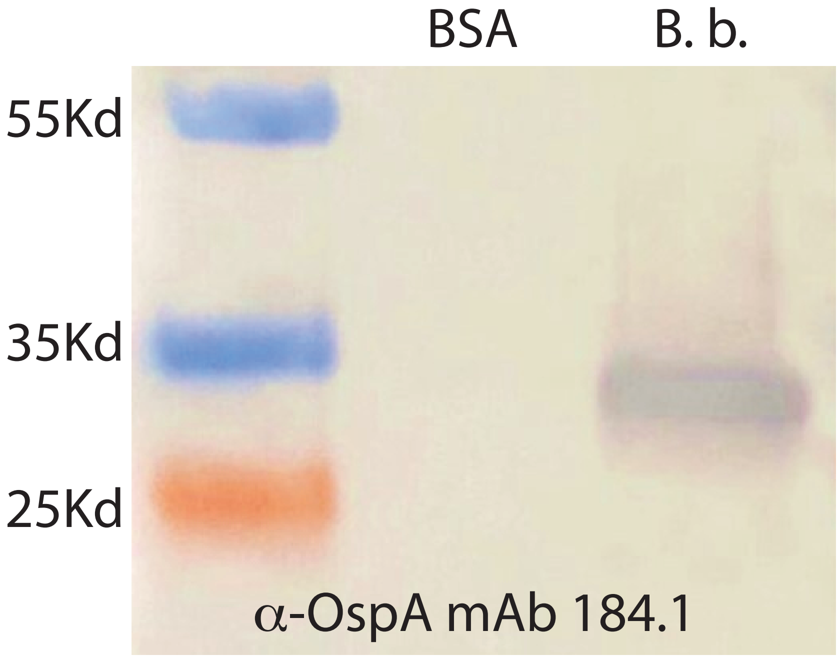


Figure S1. Cultured *B. burgdorferi* express OspA. Immunoblot of a multistrain culture of *B. burgdorferi* (B. b. passage 2 after recovery from mouse heart) against an anti-OspA monoclonal antibody. This culture was used in neutralization assays.
